# Recovery of fibroblasts from membrane attack by *S. aureus* α-toxin does not depend on acid sphingomyelinase but involves macropinocytosis

**DOI:** 10.1101/2023.12.08.570768

**Authors:** Gisela von Hoven, Claudia Neukirch, Carolin Neukirch, Martina Meyenburg, Sandra Ritz, Matthias Husmann

**Author notes:** Correspondence to: Gisela von Hoven, Matthias Husmann.

## Abstract

Damage of the plasma membrane by mechanical stress or pore forming proteins, like streptolysin O, may trigger Ca^2+^ influx-dependent repair mechanisms. Ca^2+^ influx-dependent lysosomal exocytosis, leading to release of acid sphingomyelinase, remodeling of the plasma membrane and caveolar endocytosis of membrane lesions, is reportedly involved in membrane repair after both mechanical damage or perforation by streptolysin O. Although the small β-barrel pore forming *S. aureus* α-toxin may also increase cytosolic Ca^2+^ concentration in certain cell types, and reportedly lead to release of acid sphingomyelinase from endothelial cells, evidence for a role of this response for membrane repair after attack by α-toxin is lacking. We exploited fibroblasts expressing dysfunctional acid sphingomyelinase to investigate whether this enzyme is required for membrane repair after *S. aureus* α-toxin attack. Because α-toxin-dependent loss of cellular ATP and externalization of phosphatidylserine were reversible, membrane damage by this pore former triggered an effective repair response in these cells. Although α-toxin depolarized the plasma membrane, it did not cause a simultaneous increase of [Ca^2+^]_i_. Consistently, there was no release of β-hexosaminidase, a marker of lysosomal exocytosis. Acid sphingomyelinase-deficient fibroblasts internalized α-toxin, which however did not co-localize with caveolin-1, but with FITC-dextran 70kDa, a cargo of macropinosomes. Inhibition of actin polymerization or sterol synthesis prevented recovery from α-toxin-dependent membrane damage. Therefore, we conclude that defense of fibroblasts against α-toxin does not depend on rapid calcium-influx, lysosomal exocytosis, and functional acid sphingomyelinase, but involves ongoing cholesterol synthesis and macropinocytosis.

## Introduction

Damage to the plasma membrane (PM) by physical stress or insertion of pore forming proteins (PFP) disrupts ion gradients and poses an acute threat to a cell. Because such events are inevitable during the life of an animal, evolution of membrane repair seems to be a logical consequence. One early account of membrane repair in the literature is the finding that *Xenopus laevis* eggs can efficiently reseal after damage by a glass fiber if Ca^2+^-ions are present in the surrounding milieu (Prothero et al., 1970). Cells are also able to cope with an attack by PFP, like complement (Scolding et al., 1992), perforin (Keefe et al., 2005), gasdermin (Ruhl et al., 2018) (Nozaki et al., 2022), or pore forming toxins (PFT), e.g. (Thelestam & Mollby, 1983), (Walev et al., 1994), which, due to the stable transmembrane channels they form, resist spontaneous resealing. To survive attack by PFP, target cells may remove, segregate, close, or otherwise neutralize membrane pores. However, the importance of the various mechanisms and molecular details remain a matter of debate (Babiychuk et al., 2011) (Babiychuk & Draeger, 2015) (Romero et al., 2017) (Idone et al., 2008) (Husmann et al., 2009) (Corrotte et al., 2013) (Brito et al., 2019) (Etxaniz et al., 2018) (Andrews et al., 2014) (Nygard Skalman et al., 2018). For instance, in the case of streptolysin O (SLO), a member of the cholesterol dependent cytotoxin (CDC) family of PFT, the exact contribution of microvesicle shedding (Romero et al., 2017) or endocytosis (Idone et al., 2008) to cellular recovery from attack remains unclear; possibly both pathways play a role depending on cell type and experimental conditions. As mentioned above, SLO pores or membrane lesions caused by mechanical stress, elicit a similar membrane repair response, which is initiated by rapid influx of calcium, triggering lysosomal exocytosis, release of acid sphingomyelinase (ASM), cleavage of sphingomyelin in the outer leaflet of the PM, formation of ceramid platforms, caveolar endocytosis of lesions, and in the case of SLO, proteolytic destruction of the toxin, (Andrews et al., 2014). Based on work with listeriolysin O, another CDC, Nygard et al. proposed that endocytosis might be involved in reorganization of the cell surface in response to damage, rather than in removing pores (Nygard Skalman et al., 2018). Whatever the precise function of ASM-dependent reconstitution of the damaged PM, the mechanism appears to operate in various cell types, and contribute to repair of two quite different types of membrane lesions, namely large membrane pores such as those formed by CDC, and mechanical lesions.

On the other hand, recovery from *S. aureus* α-toxin, a small β-barrel PFT (pore size ∼1.4 nm) is slow as compared to CDCs, like SLO (pore size ≥ 25 nm), (Husmann et al., 2006), which led us to propose that PFT of different structure and/or pore size trigger distinct membrane repair mechanisms. Work on other small β-PFT, including phobalysin P (PhlyP), *Vibrio cholerae* cytolysin (VCC), and aerolysin supported this idea (Gonzalez et al., 2011), (von Hoven et al., 2017), (Thapa & Keyel, 2023), (Rivas et al., 2015). Further, comparison of PhlyP and the related VCC revealed that, depending on channel width, small membrane pores can trigger, or not, Ca^2+^ influx-dependent repair mechanisms (CIDRE), (von Hoven et al., 2017). The question whether *S.aureus* α-toxin can trigger CIDRE has remained controversial. At least in some cell types, comparably high concentrations of α-toxin may cause moderate increases of cellular [Ca^2+^]_i_ (Walev et al., 1993) (von Hoven et al., 2016), but it is unclear whether this is due to ion flux through α-toxin pores of some ill defined high conductance conformation, or through cellular channel forming proteins, possibly including endogenous PFP like gasdermins, or through rupturesof the PM (von Hoven et al., 2019). A potential role of calcium-influx and ASM for membrane repair in endothelial cells treated with α-toxin was implicated by a recent report, that α-toxin induces release of this enzyme (Krones et al., 2021), but to our knowledge, there is no direct experimental evidence for ASM-dependent repair in response to α-toxin. Here we addressed the question whether ASM is *required* for membrane repair after attack by *S. aureus* α-toxin. To this end, we investigated the response of fibroblasts expressing dysfunctional ASM. This appeared sensible because normal human fibroblasts are able to recover from membrane attack by this toxin (Walev et al., 1994).

## Results

### Niemann Pick Type A disease fibroblasts (NPAF) – verification of defective ASM

Because our experimental approach relied on functionally ASM-deficient fibroblasts (NIGMS GM00112, Coriell Institute for Medical Research), we sought to confirm the defect. Using sequence analysis we found a mutation in exon 2 of the SMPD1 gene (Figure 1A), as expected (Zhang et al., 2013). Western-blot with antibodies against ASM yielded bands of equal electrophoretic mobility and similar density with extracts from normal fibroblasts (NF) and NPAF (data not shown). In order to verify that NPAF were devoid of functional ASM we measured the activity of this enzyme in cell lysates. Whereas lysates from NF reached about 17.5-fold the background level (Figure 1B), ASM activity in lysates of NPAF were hardly above buffer background. The data gave confidence that NPAF are a suitable experimental model for the investigations discussed below.

**Figure 1.**
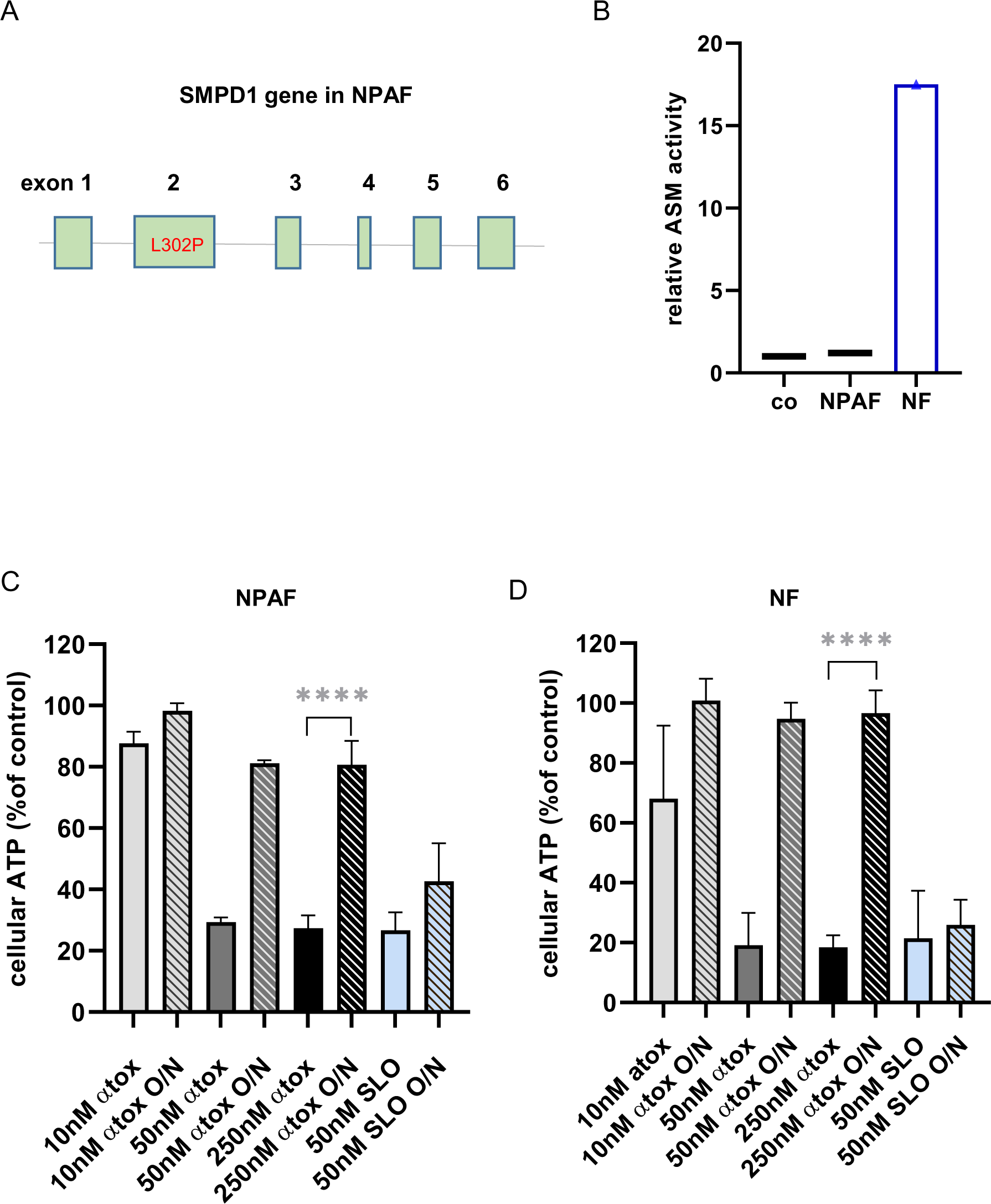
NPAF do not express functional ASM, but restore ATP-levels after loss following membrane attack by *S. aureus* α-toxin. (A): Scheme of the human SMPD1 gene. The nucleotide sequence of exons was determined in the DNA from fibroblasts (NIGMS GM00112, Niemann-Pick disease, Type A, untransformed), as detailed in the methods section. The sequence, confirming a known mutation in exon, leads to replacement of leucine by proline (indicated in red), which is common in this condition. (B) Results from ELISA for the detection of acid sphingomyelinase activity in buffer (co), or cell lysates from NPAF or NF. Mean values from two independent experiments ± SEM. (C, D) cellular ATP levels were measured in NPAF (C), or NF (D) after treatment with *S. aureus* α-toxin (α-tox), or streptolysin O-MBP (SLO) for two hours with overnight recovery in the absence of toxin (O/N), or immediately after the 2h incubation (hatched columns). Data are means from n=3 independent experiments ± SEM; significance was assessed using one-way ANOVA and Tukeýs multiple comparison test; **** denotes P< 0.0001. Samples treated with 10nM or 50nM α-toxin are from two experiments in culture media, and one in HBSS each. All of the three experiments for controls, 250nM α-toxin, or SLO were in culture media.

### *S. aureus* α-toxin induces a transient drop of cellular ATP in NPAF

Previous work showed that α-toxin at nanomolar concentrations does not permeabilize the PM, for instance of human keratinocytes, for vital dyes like trypan blue or propidium iodide (PI), but for monovalent cations, K^+^ and Na^+^ (Walev et al., 1993). Depolarization followed by loss of cellular potassium ions to the external milieu leads to a drop of cellular ATP, which can serve as a proxy for membrane damage in cells exposed to sublytic doses (i.e. doses, which do not lead to release of lactate dehydrogenase) of α-toxin. Under these conditions, human fibroblasts, HaCaT cells (keratinocyte cell line), human monocytic THP-1 cells and several other cell types may replenish major losses (> 80%) of both potassium ions and cellular ATP within a few hours.

As shown previously, normal fibroblasts are not particularly sensitive to α-toxin, but treatment of NPAF, or NF with α-toxin at 50-250nM for 2h led to a dose dependent drop of ATP down to 20% and 10%, respectively, of the untreated cells (Figure 1C and D). After overnight incubation of toxin-treated cells in toxin-free media, ATP-levels reached 95% in NF and about 80% in NPAF, indicating that dysfunctional ASM does not significantly compromise recovery from membrane damage by α-toxin.

### *S. aureus* α-toxin induces transient externalization of PS in NPAF

As a second read out for transient α-toxin-dependent perturbation of the PM in NPAF, we studied exposure of phosphatidylserine (PS) at the cell surface. In resting cells, PS is almost confined to the inner leaflet of the PM (Clarke et al., 2020). This is due to the action of flippases, enzymes located in the PM, which transport PS by an energy-dependent mechanism against a gradient to the interior leaflet of the PM. Scramblases are also located in the PM, but these enzymes can rapidly move lipids in either direction. Exposure of PS at the exterior side may result from various kinds of cellular stress and typically precedes apoptosis or necroptosis. In the presence of Ca^2+^ ions, annexin-V, or a labelled derivative, binds to PS exposed at the cell surface, where it can be conveniently detected, for instance by fluorescence-activated cell sorting (FACS). Reportedly, treatment of A549 cells with 2.5µg/ml *S. aureus* α-toxin for 2h led to ∼75% increase of annexin V binding (Lizak & Yarovinsky, 2012). In NPAF, treatment with α-toxin, (250nM - corresponding to ∼ 8μg/ml - for 2h at 37°C in HBSS), roughly doubled annexin-V-binding to cells (Figure 2A and B). An oligomerization-deficient mutant of α-toxin (H35R) did not cause a significant increase (Figure 2B), suggesting that α-toxin-dependent PS exposure requires oligomerization, and presumably pore formation. As compared with NPAF, a smaller population of toxin-treated NF bound labelled annexin-V, but recovery in the absence of toxin reduced the fraction of annexin-V-binding cells by 50%, as in NPAF (not shown). Fibroblasts are heterogeneous and NPAF are probably not coisogenic with the NF used here. Moreover, culture conditions of NF and NPAF are different, precluding meaningful comparison. At any rate, the present data obtained with NPAF documents that cells devoid of functional ASM are able to restore normal PS-distribution between leaflets of the PM and replenish intracellular levels of ATP.

**Figure 2.**
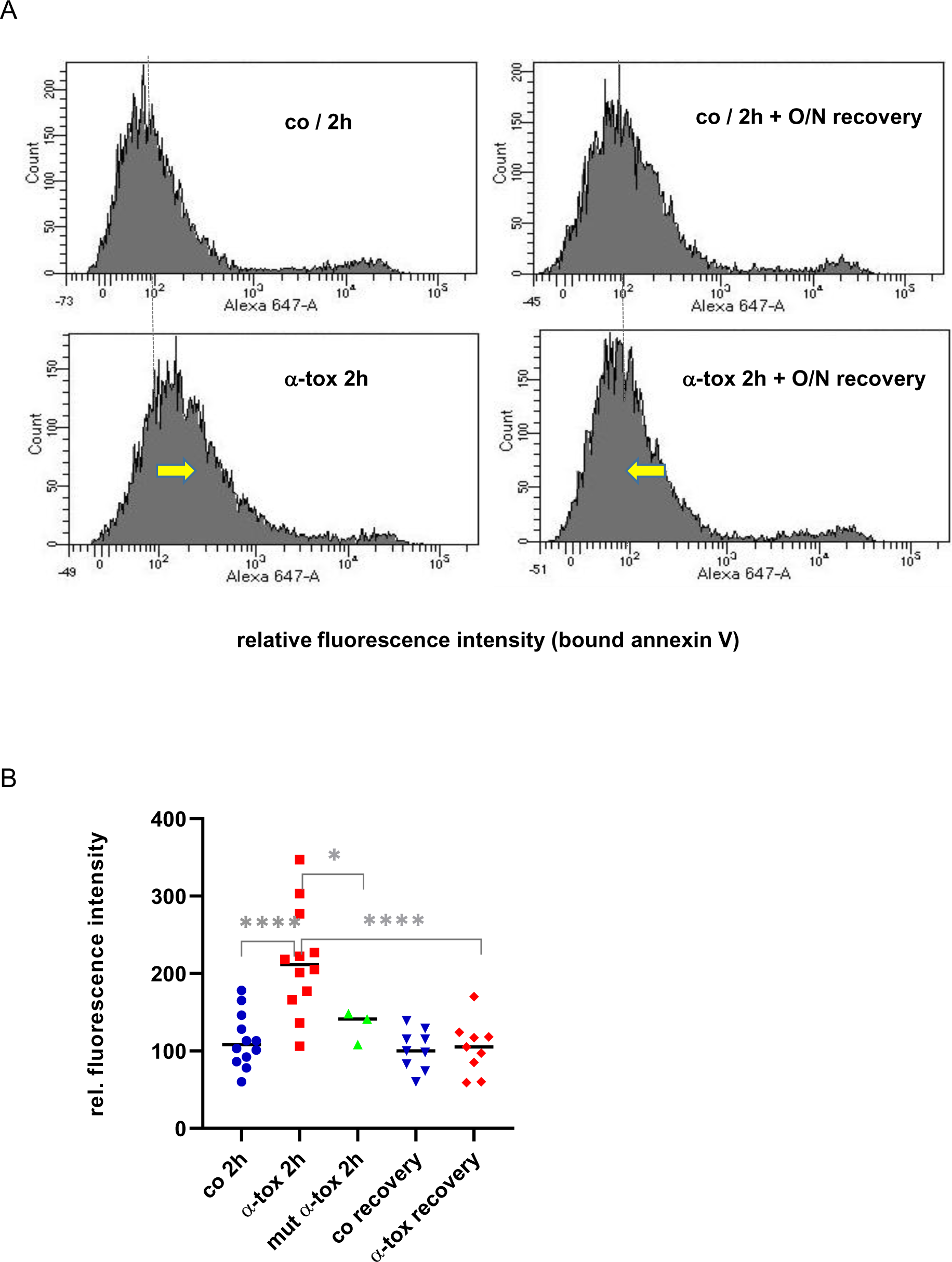
*S. aureus α*-toxin causes transient exposure of PS on NPAF. (A) FACS-analysis of Alexa647-annexin V-binding (x-axis) to NPAF treated for 2h with *S. aureus* α-toxin (250nM), (lower left panel), or after removing toxin and O/N recovery (lower right panel). There is an increase of fluorescence intensity at 2h, and a reversal of this increase after recovery, as indicated by yellow arrows. Upper panels show corresponding data from untreated controls (co) incubated for 2h only (left), or 2h plus O/N incubation after washing (right). (B) Graph summarizing multiple FACS experiments as shown in (A) with an additional set of data obtained with pore-dead mutant α-toxin H35R (green symbols). Relative fluorescence intensities (median channels) are from n ≥ 3 experiments for each treatment; colored symbols indicate values of individual experiments, horizontal black lines indicate medians. Toxin-dependent increase of annexin V-binding and subsequent decrease following O/N recovery (red symbols) were highly significant (* and **** denote P<0.0001**** and P<0.05 respectively, as determined by one-way ANOVA and Tukeýs multiple comparison).

### *S. aureus* α-toxin does not trigger rapid changes of [Ca^2+^]_i_ in NPAF

Influx of Ca^2+^ is considered the principle trigger of PM repair (Cooper & McNeil, 2015). *S. aureus* α-toxin may increase [Ca^2+^]_i._ in certain cell types, for instance lung epithelial cells (Eichstaedt et al., 2009), keratinocytes (HaCaT cells), (von Hoven et al., 2016), and endothelial cells (Krones et al., 2021). Yet, it is not clear whether these increases are due to Ca^2+^ influx through toxin pores, other channels, or some secondary lesions caused by endogenous PFP, or through ruptures. For triggering CIDRE, it appears to be necessary that Ca^2+^ influx occurs through the pore to be repaired. Thus, SLO or PhlyP, which elicit CIDRE, do not afford protection against simultaneous challenge with α-toxin (Husmann et al., 2006), or VCC (von Hoven et al., 2017), respectively, suggesting that even robust calcium-influx will only trigger repair of the membrane pore, which causes the influx. So far, there is no evidence that α-toxin-dependent increases of [Ca^2+^]_i._ can trigger productive CIDRE. Here we compared the response of NPAF loaded with Fluo-8, a calcium-sensitive dye, to α-toxin, or SLO, a *bona fide* inducer of calcium-influx. First, we performed experiments in a plate reader format to detect potential toxin-dependent changes in the entire cell population. Exposure of NPAF to SLO induced a sharp increase of fluorescence intensity, but there was no rise with equimolar concentrations of α-toxin (Figure 3A). PhlyP, a small β-PFT forming larger pores than α-toxin caused increases of [Ca^2+^]_i._, in line with our previous results (von Hoven et al., 2017). To observe responses in single cells, we cultured NPAF in flow-chambers, loaded them with Fluo-8, and analyzed them by video microscopy under cell culture conditions. Although injection of SLO induced a rapid and strong increase of fluorescence, we did not discern any increase in α-toxin-treated cells (see images in Figure 3B).

**Figure 3.**
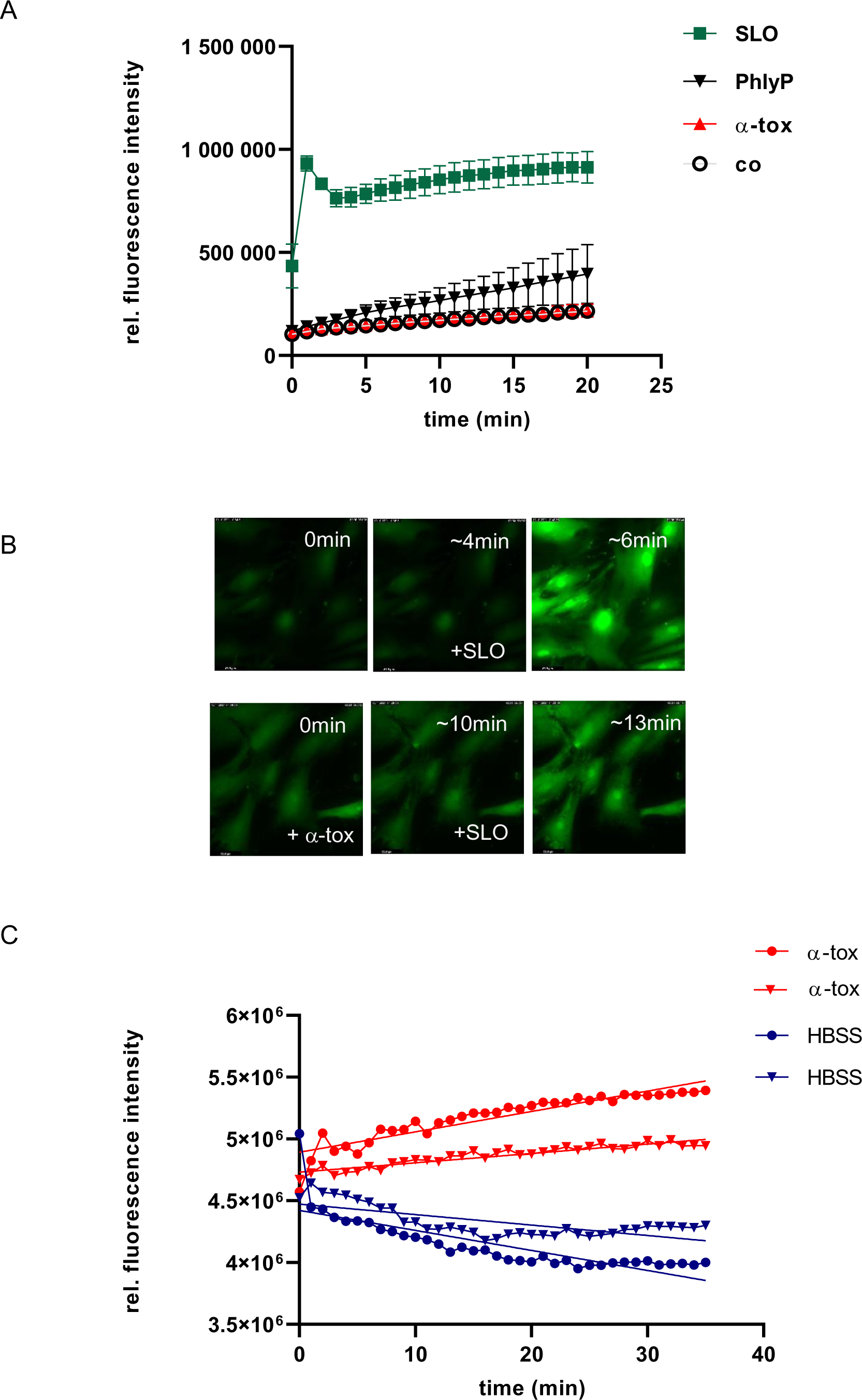
*S. aureus α*-toxin does not alter [Ca] in NPA-fibroblasts, although it depolarizes the PM. (A) Kinetics of fluorescence in Fluo-8-loaded NPAF culture, treated with SLO, PhlyP or α-toxin. “co” denotes treatment with buffer. Mean values of n=3 independent experiments ± SEM. Differences of fluorescence intensities at 20min were highly significant (P<0.0001) for the comparison of buffer *vs.* SLO or PhlyP, but not significant for buffer vs. α-toxin, as determined by one-way ANOVA and Tukeýs post hoc test. (B) Sequential images from video microscopy of Fluo-8AM-loaded NPAF. Upper panels (positive control): after short incubation without toxin, SLO (50nM) was added; note rapid increase of fluorescence. Lower panels: after short incubation without toxin, α-toxin (250nM) was added, but no significant change of fluorescence was noted during the next 10min; subsequent addition of SLO led to rapid increase of fluorescence intensity, although to lower extent as compared to the positive control. (C) Kinetics of fluorescence in DISBAC2(3)-loaded NPAF culture treated with α-toxin or buffer (HBSS); data from two out of three similar, independent experiments. Identical shape of symbols indicates that data points belong to the same experiment; same color indicates same treatment. Simple linear regression analysis revealed highly significant (P< 0.0001) deviation from 0-slope for both, HBSS (negative slope) and α-toxin (positive slope).

### *S. aureus* α-toxin rapidly depolarizes the plasma membrane of NPAF

Because α-toxin did not appear to cause rapid elevation of [Ca^2+^]_i_ in NPAF, we sought to confirm that within the period of observation α-toxin altered membrane integrity. For this purpose, we used DiSBAC2(3), a fluorescent slow response reporter of PM potential. We found that α-toxin increases DiSBAC2(3)-fluorescence in NPAF in the same time frame where it fails to increase fluorescence of Fluo-8 (Figure 3C). Signal strength and kinetics obtained with DiSBAC2(3) were similar as previously observed with α-toxin-treated airway epithelial cells (Eiffler et al., 2016). Taken together our above results from Fluo-8 based assays and those obtained with DiSBAC2(3) suggests that α-toxin depolarizes the PM of NPAF without significantly changing overall [Ca^2+^]_i_ in the same time frame.

### *S. aureus* α-toxin does not trigger significant release of β-hexosaminidase from NPAF

Upon wounding of the PM, or perforation by a PFT like SLO, the resulting influx of Ca^2+^ ions triggers lysosomal exocytosis and consequent externalization of ASM and other lysosomal content like β-hexosaminidase. Here, we measured the activity of β-hexosaminidase in cultures of SLO- or α-toxin-treated NPAF. SLO induced rapid and robust increases of β-hexosaminidase activity, as expected (Figure 4A). In contrast, cultures receiving α-toxin remained negative, suggesting that no significant lysosomal exocytosis occurred.

**Figure 4.**
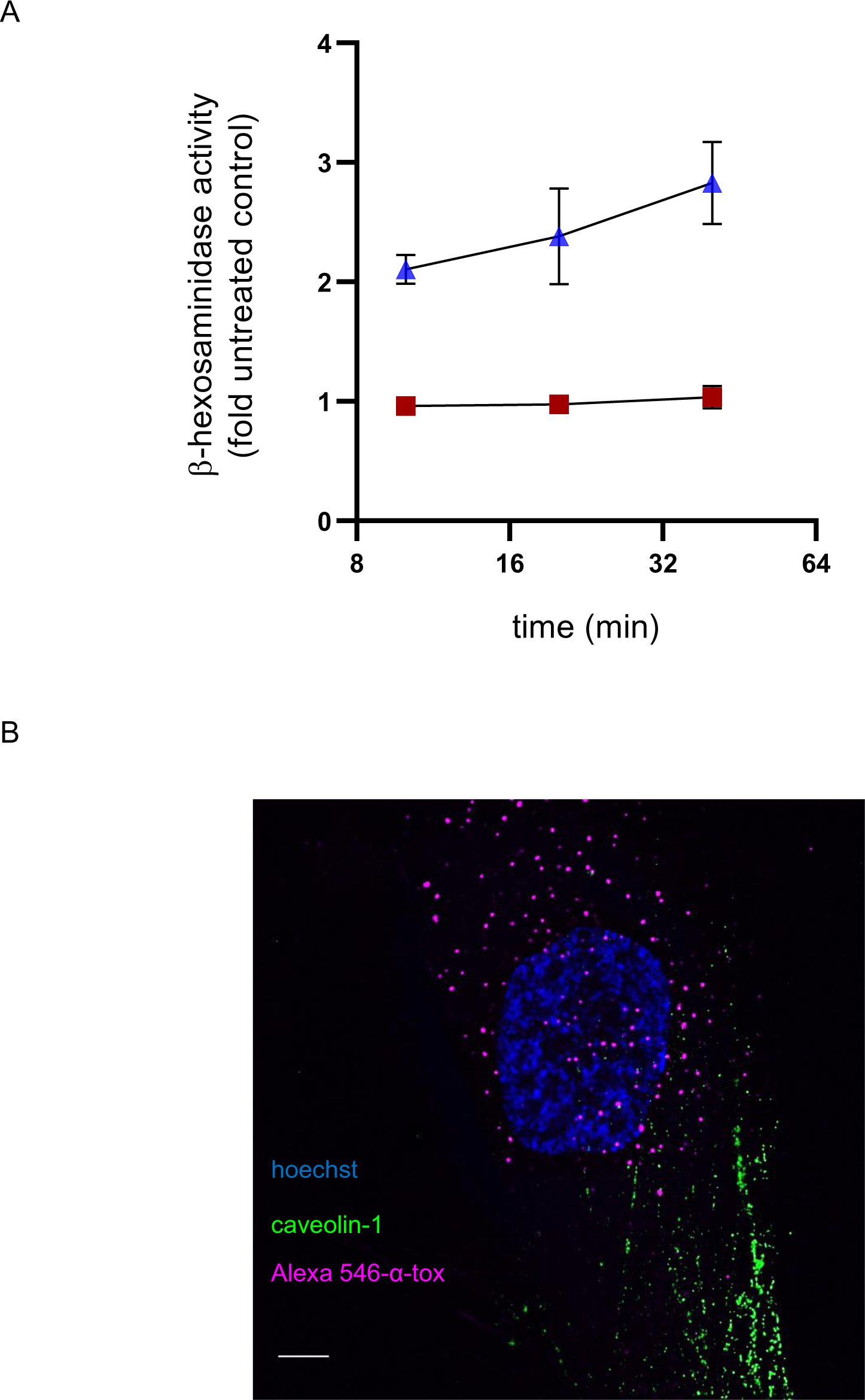
*S. aureus α*-toxin does not trigger lysosomal exocytosis in NPAF. No co-localization with caveolin-1. (A) Kinetics of β-hexosaminidase release from NPAF culture, treated with SLO (blue symbols), or α-toxin (red). Mean values of n=3 independent experiments ± SEM. Simple linear regression revealed significant (P=0.0179) deviation from 0-slope for SLO, but no significance for α-toxin (P=0.139). (B) Representative IF microscopic image (wide field, deconvolution), of an NPAF cell, treated with Alexa546-α-toxin for 15 min and stained for caveolin-1. Scale bar, 5 μm.

### Evidence for uptake of *S. aureus* α-toxin by NPAF *via* macropinocytosis

Reportedly, ASM-dependent membrane repair of SLO-pores involves enzymatic remodeling of the outer leaflet of the PM with the formation of local ceramide platforms, followed by caveolar internalization of membrane pores (Andrews et al., 2014). Previously we have found that epithelial cells internalize *S. aureus* α-toxin, whereas other cell types (e.g. Huh7, a liver carcinoma cell, expressing very low levels of caveolin) fail to do so. Importantly, the ability to internalize α-toxin correlated with the ability of cells to cope with membrane damage by this PFT (Husmann et al., 2009). To investigate whether NPAF too can internalize α-toxin, we first analyzed samples treated with Alexa546-labelled α-toxin in a double staining approach on samples either permeabilized with the detergent triton-X100, or not. In non-permeabilized cells, there were many puncta corresponding to Alexa546-labelled α-toxin, which however escaped immunostaining with antibody. Because Alexa546-labelled α-toxin in permeabilized samples became accessible to immunostaining, puncta became double positive (supplementary Figure 1). This experiment showed that NPAF do internalize α-toxin. However, we did not observe co-localization of Alexa546-labelled α-toxin and caveolin-1 in a two-color fluorescence experiment (Figure 4B).

There is preliminary evidence, based on the use of inhibitors that epithelial cells internalize α-toxin via macropinocytosis (Shah et al., 2018), (von Hoven & Husmann, 2019). In order to corroborate the available data and extend the investigation to NPAF, we co-incubated these cells with Alexa546-labelled α-toxin and FITC-dextran 70kDa, which is too large to enter cells via clathrin coated pits or caveolae (Canton, 2018). Under the fluorescence microscope, we observed numerous double-labelled puncta (Figure 5, panels a through d), most of which had diameters of > 300 nm, compatible with macropinosomes (Canton, 2018), (Kerr & Teasdale, 2009). Importantly, results in HaCaT cells, where uptake of α-toxin was first demonstrated (Husmann et al., 2009), were similar (supplementary Figure 2). Macropinocytosis is an actin dependent process (Canton, 2018), (Kerr & Teasdale, 2009). Treatment with 20µM cytochalasin D (CYD), which inhibits actin-polymerization, altered the shape of FITC-dextran 70kDa-positive puncta and appeared to modulate uptake of Alexa546-labelled α-toxin (Figure 5, panel e through h). Size and shape of FITC-dextran 70kDa-positive puncta in controls appeared more homogenous than in CYD treated samples. Whereas Alexa546-labelled α-toxin in controls formed distinct puncta, much of the Alexa546-staining in CYD treated cells appeared diffuse with fewer Alexa546-positive puncta co-localizing with FITC-dextran 70kDa.

**Figure 5.**
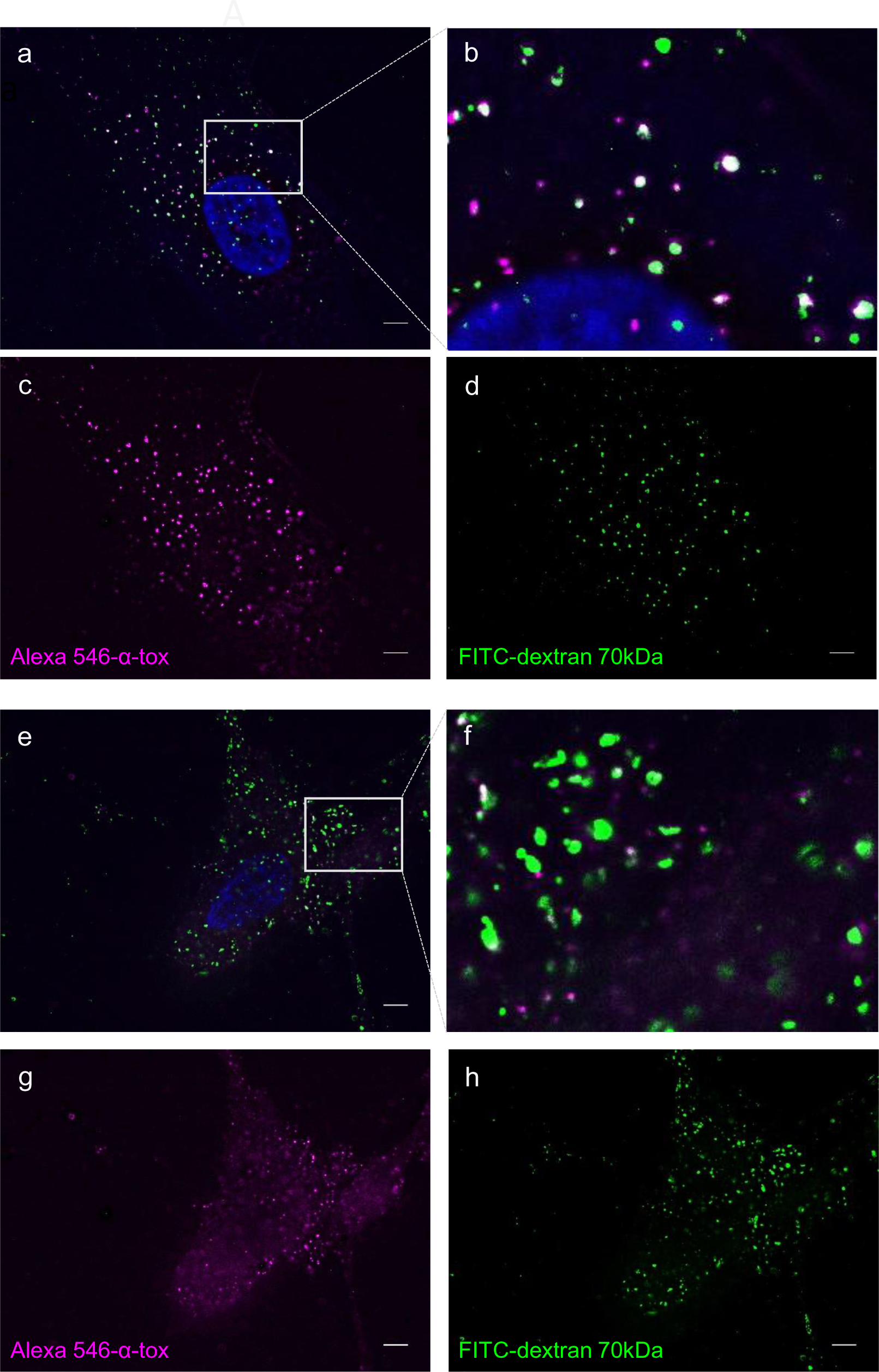
*S. aureus α*-toxin co-localizes with FITC-dextran 70kDa in NPAF. Cytochalasin D modulates uptake. Representative IF microscopic image (wide field, deconvolution), of an NPAF cell, simultaneously treated with Alexa546-α-toxin and FITC-dextran 70kDa. Samples corresponding to panels *a* through *d* received solvent, panels *e* through *h* 20µM CYD. Scale bars, 5 μm.

### Cytochalasin D prevents recovery of NPAF after attack by *S. aureus* α-toxin

If macropinocytosis was important for survival of NPAF exposed to α-toxin, CYD should prevent recovery after attack by the PFT. In fact, this drug inhibits normalization of ATP-levels in α-toxin-treated NF (Valeva et al., 2000). Therefore, we analyzed its effect on annexin-V binding and influx of PI in NPAF (Figure 6). Incubation of NPAF with α-toxin led to transient increase of staining with annexin-V, as expected from the experiment shown in Fig.2. Incubation with CYD too caused a moderate increase of annexin V-binding, which was not reversible. However, the combination of CYD and α-toxin during the 2h-incubation prevented subsequent recovery from membrane attack, as it enhanced binding of Alexa647-annexin-V and increased influx of PI (Figure 6A and B).

**Figure 6.**
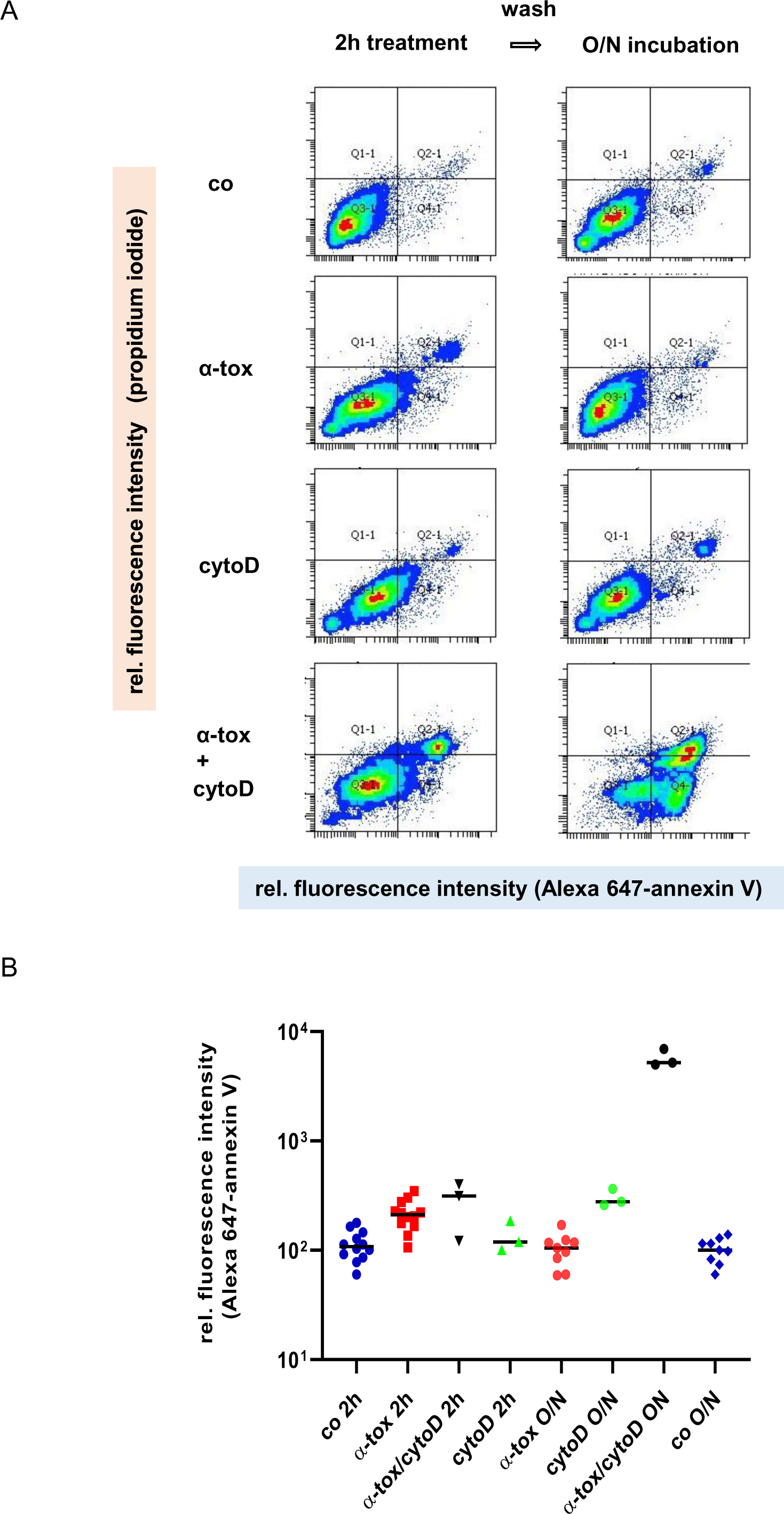
Cytochalasin D enhances *S. aureus α*-toxin dependent damage of NPAF. (A) FACS-analysis of NPAF treated, or not, with α-toxin, and/or 20µM CYD, or solvent (co). Left set of panels: treatment for 2h before staining with Alexa647-annexin V and PI. Right: treated as in corresponding left panels, but subsequently washed and incubated O/N before staining with Alexa647-annexin V and PI. (B) Summary of Alexa647-annexin V binding data from ≥ 3 experiments as shown in (A). Note log scale. Data points (rel. fluorescence intensity) are median channels from individual experiments with n ≥ 3 independent experiments for each treatment; short, horizontal lines indicate medians. Annexin V-binding following O/N incubation with α-toxin plus 20µM CYD was significantly higher compared to all other samples (P<0.0001, one-way ANOVA and Tukeýs multiple comparison).

### Inhibitors of cholesterol synthesis inhibit cellular recovery from α-toxin attack

Among the most efficient inhibitors of ATP-replenishment in NF after α-toxin-attack is cerulenin (Valeva et al., 2000), an inhibitor of fatty acid-an sterol-synthesis (Nomura et al., 1972). Macropinocytosis is subject to regulation by cholesterol (Kerr & Teasdale, 2009), and the sterol regulatory element binding protein, SREBP-2, is implicated in membrane repair (Gurcel et al., 2006). Therefore, we investigated the effect of triparanol (TRP), a selective inhibitor of cholesterol synthesis (Estes, 1960), (for a simplified scheme see Figure 7A). Treatment with TRP increased toxin-dependent annexin V binding already during the 2h period of co-incubation with α-toxin, suggesting that cellular cholesterol synthesis is of vital importance for NPAF early during an attack by α-toxin (Figure 7B). As illustrated in Figure 7A, TRP causes accumulation of desmosterol, a ligand of liver X receptor (Yang et al., 2006). This prompted the question whether the effect of TRP in our experiments might be due to desmosterol. However, exogenous desmosterol did not mimic the effect of TRP (Figure 7C), and other inhibitors of cholesterol synthesis, AW9944 and lovastatin, which do not lead to accumulation of desmosterol, caused increased binding of annexin V as well (data not shown). These results indicate that cholesterol synthesis is important for survival of NPAF attacked by α-toxin.

**Figure 7.**
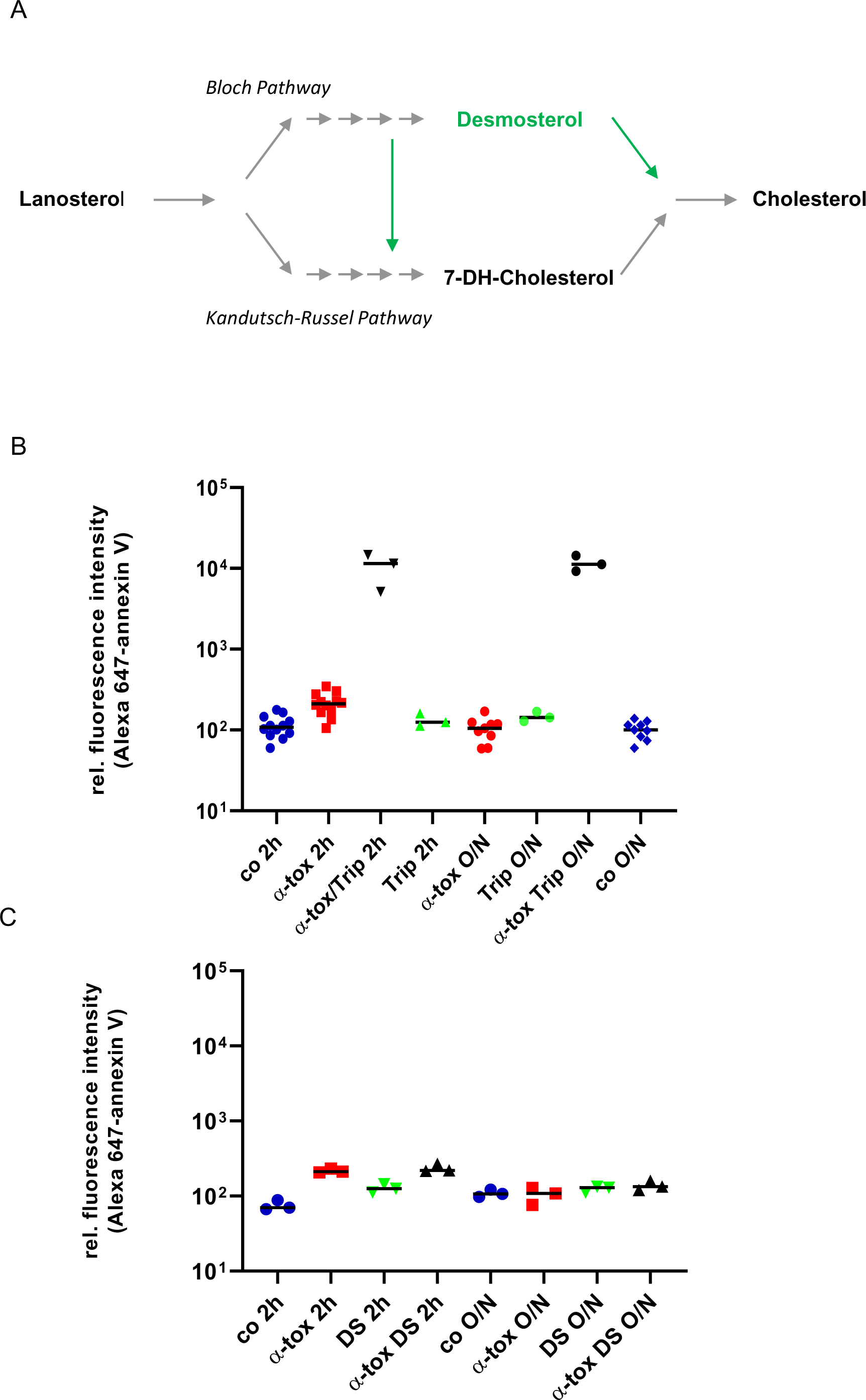
Triparanol enhances *S. aureus α*-toxin dependent damage of NPAF. (A) Simplified scheme of cholesterol synthesis downstream of lanosterol, highlighting the bifurcation of the synthetic pathway. TRP inhibits steps symbolized by green arrows, and causes accumulation of desmosterol. (B) NPAF were treated, or not, with α-toxin in the presence of 45µM TRP, or solvent. After 2 hours, or wash followed by O/N recovery in media without TRP, FACS-analysis was performed to measure binding of Alexa647-annexin V and staining with PI. The graph shows Alexa647-annexin V binding data from n ≥ 3 independent experiments. Note log scale. Short, horizontal lines indicate medians. Annexin V-binding after incubation with α-toxin plus 45µM TRP (black symbols) was far higher than in all other samples (P<0.0001, one-way ANOVA and multiple comparison with Tukeýs post hoc test). Note strong increase of annexin V-binding already after 2h α-toxin plus 45µM TRP. (C) NPAF were treated, or not, with α-toxin in the presence of 10µM swamosterol (DS), or solvent alone. After 2 hours, or wash followed by O/N recovery in media without DS, FACS-analysis was performed to measure binding of Alexa647-annexin V and staining with PI. The graph shows data on the binding of Alexa647-annexin V from n ≥ 3 independent experiments. Log scale. Short, horizontal lines indicate medians. Annexin V-binding after incubation with α-toxin plus DS (black symbols) was not significantly higher than in samples receiving α-toxin plus solvent. (one-way ANOVA and multiple comparison with Tukeýs post hoc test).

## Discussion

A series of previous publications implicated ASM in membrane repair after damage by the pore forming bacterial toxin SLO or mechanical stress, suggesting broad relevance of this mechanism for mending various types of membrane lesions (Andrews et al., 2014). On the other hand, there is evidence, that membrane repair after damage by *S. aureus* α-toxin depends on different mechanisms (Husmann et al., 2009) (Husmann et al., 2006), (von Hoven et al., 2019), (Valeva et al., 2000). However, according to a recent report, α-toxin may trigger release of ASM from cells (Krones et al., 2021), although the functional significance of this finding is not known. In order to clarify whether ASM is a *requirement* for membrane repair after attack by α-toxin we decided to investigate fibroblasts lacking functional ASM. We verified the underlying mutation in the SMPD1 gene and exploited these fibroblasts (NPAF) to show that cells do not require functional ASM for recovery from an attack by *S. aureus* α-toxin.

Experimental evidence for α-toxin-dependent transient perturbation of the PM in NPAF was two-fold. First, cells replenished intracellular ATP levels after a massive α-toxin-induced drop. Second, transient increases of annexin V binding indicated reversible exposure of PS at the cell surface. It is possible that externalization of PS is secondary to α-toxin-dependent loss of ATP, because flippases, which move PS to the interior leaflet of the PM depend on energy. Lipid scramblases could also mediate α-toxin-induced exposure of PS, but these enzymes may also be associated with membrane repair. Thus, scramblase TMEM16F assumes a role in PS exposure and membrane repair following perforation by CDC (Wu et al., 2020); and phospholipid scramblase 1 confers IFN type I-dependent cellular resistance to α-toxin (Lizak & Yarovinsky, 2012), (Yarovinsky et al., 2008). That NPAF can reverse both toxin-dependent externalization of PS and a major drop of cellular ATP, indicates that ASM is not essential for reconstitution of PM integrity after damage by α-toxin.

In addition to ASM, lysosomal proteases contribute to the remodeling of damaged PM (Castro-Gomes et al., 2016). Because α-toxin did not lead to lysosomal exocytosis in NPAF, lysosomal proteases are unlikely to play a role in this context. Reportedly, ASM-dependent membrane repair involves endocytosis of membrane pores by caveolae (Andrews et al., 2014). Although internalized by NPAF, AlexaFluor546-labelled α-toxin did not co-localize with caveolin-1 in our experiments, not excluding though co-localization at other time points. *S. aureus* α-toxin, in contrast to SLO, did not induce a rapid rise of [Ca^2+^]_i_, in fibroblasts, although it depolarized the PM. Together this suggests that neither rapid Ca^2+^ influx, nor downstream events, including ASM-dependent remodeling of the PM are critically involved in membrane repair after exposure to α-toxin. In order to survive membrane damage by α-toxin, NPAF must be in command of different rescue mechanisms.

Previously we have found that immortalized human keratinocytes, HaCaT, internalize α-toxin monomers and oligomers, a process closely followed by replenishment of cellular ATP (Husmann et al., 2009). HaCaT cells co-internalize fluorescently labelled toxin with cross-linked GPI-anchored proteins (von Hoven et al., 2019), and uptake of the PFT involves non-canonical functions of regulators of translation (Kloft et al., 2012), (von Hoven et al., 2019). Inhibition of toxin-internalization is paralleled by inhibition of cellular recovery (Husmann et al., 2009), (Kloft et al., 2012). Because amiloride and cytochalasin D (CYD) inhibited endocytosis of α-toxin by epithelial cells, a potential role of macropinocytosis (Canton, 2018), (Kerr & Teasdale, 2009) has been discussed in this context (Shah et al., 2018), (von Hoven & Husmann, 2019). The present data revealing co-localization of fluorescent-labelled α-toxin and FITC-dextran 70kDa in large endosomes of both NPAF and HaCaT cells supports this idea. Further, CYD modulates recovery of NPAF from α-toxin-dependent membrane damage, in line with previous findings in NF (Valeva et al., 2000). Taken together, these results suggest that fibroblasts and epithelial cells utilize macropinocytosis to internalize α-toxin and thereby survive membrane perforation by this PFT. Its capacity to rapidly engulf large areas of PM (Watanabe & Boucrot, 2017), would indeed seem to predestine macropinocytosis as an efficient defense mechanism against PFP.

Although occurring constitutively, macropinocytosis is subject to regulation. Intriguingly, an endogenous PFP drives macropinocytosis in the toad *B. maxima* (Zhao et al., 2022). In mammalian cells, several signaling proteins associated with macropinocytosis become activated upon membrane damage by α-toxin, for instance the epidermal growth factor receptor EGFR (Palm, 2019), (Haugwitz et al., 2006). In dendritic cells, latex beads or dextran 70kDa strongly increase the rate of macropinocytosis by inducing a slow rise of [Ca^2+^]_i._ (Falcone et al., 2006), similar to what one can observe in some cell types treated with α-toxin. Like the formation of macropinosomes (Palm, 2019), cellular recovery from α-toxin is sensitive to inhibitors of PI3K (von Hoven et al., 2015), (Kloft et al., 2010). Macropinocytosis, an important mechanism of nutrient uptake becomes activated upon cellular starvation (Palm, 2019). Because membrane damage by α-toxin causes cellular starvation, it could promote macropinocytosis. The PFT causes a drop of ATP and inhibits cellular uptake of leucine, thereby activating AMP-activated protein kinase (AMPK) and amino acid starvation sensor GCN2 (general control non-derepressable 2), respectively (von Hoven et al., 2015), (Kloft et al., 2010), (Kloft et al., 2012). It is known that AMPK promotes macropinocytosis (Kim et al., 2018) and autophagy, two adaptive responses to starvation (Florey & Overholtzer, 2019), which may be both triggered by α-toxin, (Kloft et al., 2010), and the present work. Although GCN2 does not seem to activate macropinocytosis *per se*, it adapts cellular physiology to scavenging dependent growth (i.e. growth based on the use of extracellular protein as an amino acid source). Activated GCN2 phosphorylates eIF2α (Kloft et al., 2010). In epithelial cells treated with α-toxin, p-eIF2α accumulates next to membrane bound toxin where it recruits PPP1r15B (Kloft et al., 2012), the constitutive regulatory subunit of eIF2α-phosphatase. Through its N-terminal amphipathic α-helix, PPP1r15B alters membrane curvature, providing a mechanism for endocytosis of α-toxin (Kloft et al., 2012). Because PPP1r15B interacts with G-actin (Chen et al., 2015), it is apt to integrate conserved stress signaling in response to membrane damage and cytoskeletal rearrangements relevant to repair. Further investigation is required to elucidate a potential role of PPP1r15B/G-actin interaction for uptake of α-toxin by macropinocytosis.

A characteristic property of macropinocytosis is its bi-directionality (Falcone et al., 2006), as exemplified by prion-like proteins, which enter cells via macropinocytosis and exit with intraluminal vesicles (ILV) via back fusion of multivesicular bodies with the PM (Bloomfield & Kay, 2016). Taken together, previous results (Husmann et al., 2009) and the present data suggest that the stabile α-toxin pore might take the same route. Concerns have been expressed that PFT pores could continue to perturb cellular ion concentrations after endocytosis, raising doubts that endocytosis could lead to their neutralization (Keyel et al., 2011). However, segregation of membrane pores in ILV may curb their effects on cellular ion physiology, and also provide a way to concentrate pores for subsequent exocytosis, thus minimizing the expenditure of membrane lipids. In this regard, the apparent role of sterol synthesis for cellular recovery from attack by α-toxin (this work) or aerolysin (Gurcel et al., 2006) might reflect the increased membrane turnover associated with macropinocytosis and exocytosis of PFT. Reportedly, LC3-associated macropinocytosis is important for the restructuring of the PM after LASER-induced damage and rapid CIDRE type responses (Sonder et al., 2021). Based on the collective data we believe that influx (if any) of Ca^2+^ ions through α-toxin pores in the PM of fibroblasts – and possibly any target cell - is insufficient to trigger CIDRE. While patch repair (Thapa & Keyel, 2023), or covalent modification of pore complexes, for instance by ubiquitination, as recently discussed (von Hoven et al., 2022), may contribute to repair of those small β-PFT, which form channels of low conductance for Ca^2+^, macropinocytosis emerges as a candidate mechanism, which could accomplish timely removal of such pores and concomitant structural restoration of the membrane.

To sum, fibroblasts devoid of functional acid sphingomyelinase (ASM) are able to recover from an attack by *S. aureus* α-toxin. As summarized in Figure 8, the small β-PFT depolarizes the PM, causes energy loss and exposure of phosphatidylserine (PS), but it does not trigger rapid influx of Ca^2+^. Therefore, calcium-influx dependent repair mechanisms are unlikely to play a role in this experimental model. We propose that conserved mechanisms of nutrient-sensing and -traffic, including macropinocytosis, are employed to recognize membrane damage, engulf α-toxin, and cope with metabolic consequences of membrane perforation by this small β-PFT.

**Figure 8.**
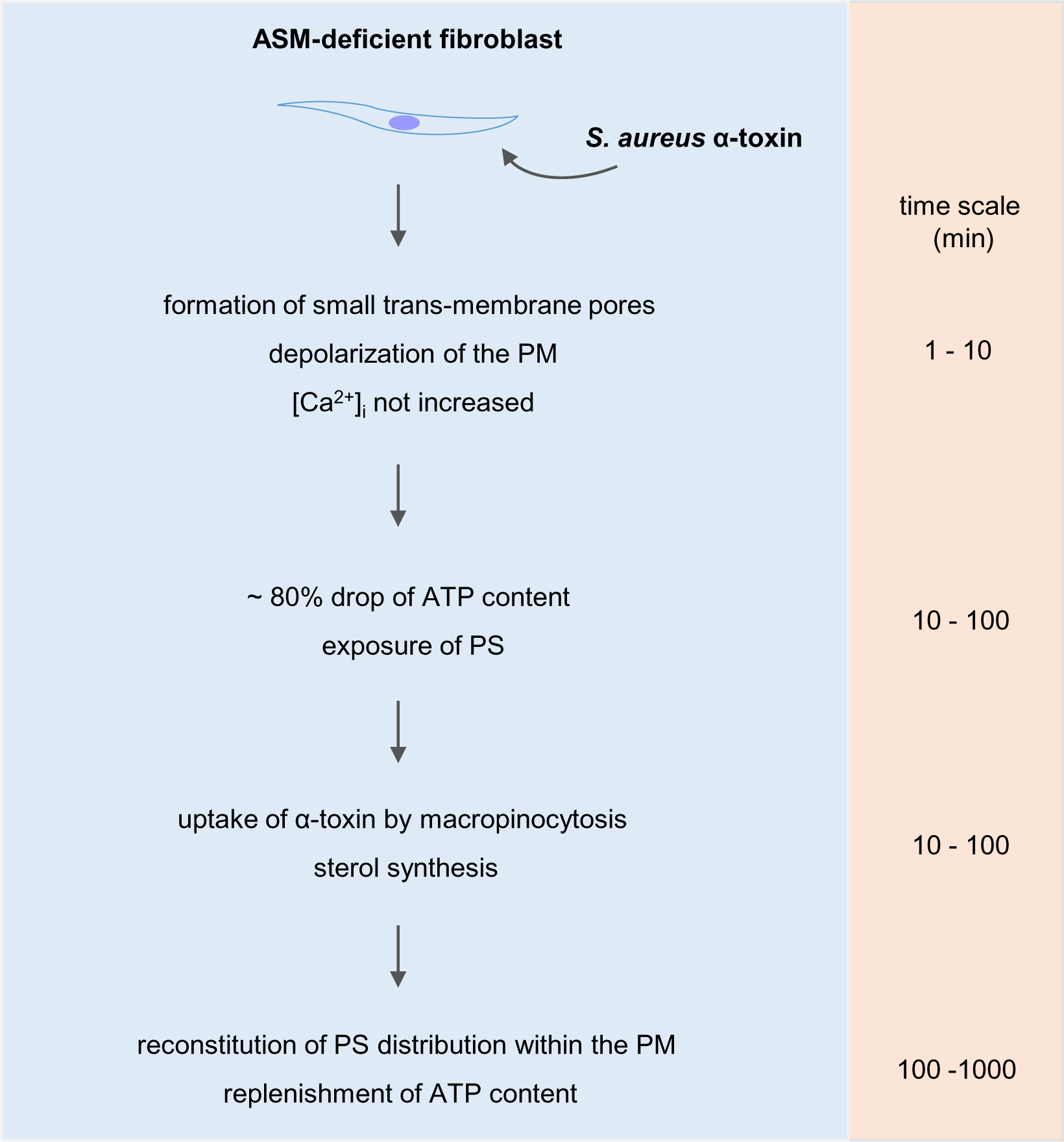
ASM is not required for membrane repair after attack by *S. aureus* α-toxin. Fibroblasts lacking functional acid sphingomyelinase (ASM), an enzyme implicated in membrane repair/remodeling following attack by the large pore forming streptolysin O (pore diameter ca. 30 nm), are able to recover from small pore forming α-toxin (pore diameter ca. 1.4 nm), similar to normal fibroblasts (Walev et al. 1994). The α-toxin pore depolarizes the PM, leading to a drop of cellular ATP content, and exposure of phosphatidylserine (PS), but it does not trigger rapid calcium-influx. Conserved mechanisms of nutrient sensing and traffic, including macropinocytosis, along with sterol synthesis, are important to overcome membrane damage by α-toxin.

## Materials and Methods

### Antibodies

Rabbit polyclonal anti-human caveolin-1 (N20), and FITC-dextran 70kDa were from Santa Cruz Biotechnology Inc. (cat # c-894 and cat # sc-263323, respectively); goat polyclonal anti-rabbit IgG-Alexa Fluor^TM^ 488 conjugate (A-11034) and annexin V Alexa Fluor^TM^ 647 (cat # A23204) were from Thermo Fisher Scientific. Rabbit polyclonal anti-*S. aureus* α-toxin was from Sigma-Aldrich (cat # S7531); F(ab)_2_ fragments were produced and purified by standard procedures.

### Chemicals

Amiloride hydrochloride, Sigma (A7410); AY9944, Santa Cruz Biotechnology Inc. cat # sc-202965; CalciFluor^TM^ Fluo-8AM was from Santa Cruz Biotechnology Inc, cat # sc-362561); Cremophor®, Millipore (cat # 23840); Cytochalasin D, Sigma (C8273); Desmosterol, Avanti (cat # IO-005808.2); DiSBAC2(3), Thermo Fisher Scientific (cat # B413); Lovastatin, ChemCruz (cat # sc-200850); 4-Methylumbelliferyl-N-Acetyl-ß-D-glucosaminide, Sigma (M2133); Paraformaldehyde, Roth; Phenylmethylsulfonyl fluoride (PMSF), Sigma-Aldrich (P7626); Probenecid, Sigma (P8761); ProLong^TM^ Diamond Antifade Mountant, (Invitrogen P36965); Triparanol, ChemCruz (cat # sc-208472).

### Toxins

*S. aureus* α-toxin, constitutively active streptolysin O (C530A)-maltose binding fusion protein (SLO-MBP), and pro-phobalysin P, (pPhlyP), were made and purified as described earlier (Jursch et al., 1994), (Weller et al., 1996), (Rivas et al., 2015). For brevity, streptolysin O-MBP or pPhlyP are termed SLO and PhlyP, respectively, in the text.

### Cells and culture conditions

NPAF, i.e. Niemann-Pick Disease Type A untransformed fibroblast cells (NIGMS GM00112), were obtained from the Coriell Institute for Medical Research, Camden, NJ, U.S.A.. Normal human fibroblasts (NF) were a kind gift from Professor B. Plachter (Institute of Virology at the University Medical Center, Johannes Gutenberg-Universität Mainz, Germany). NF were generally cultured in DMEM/F-12 with Glutamax (Ref. 31331-028) with 5% heat-inactivated FCS, penicillin/streptomycin (Sigma), 10mM HEPES (Sigma). NPAF were cultured in DMEM/F-12 with Glutamax (Ref. 31331-028) with 10% non-heat-inactivated FCS, 1% non-essential amino acids (GIBCO), penicillin/streptomycin (Sigma), 10mM HEPES (Sigma). The human keratinocyte cell line HaCaT was cultured as described (Haugwitz et al., 2006). All media and medium additives were from Gibco/Life Technologies unless stated otherwise.

### Sequence analysis of SMPD1

From genomic DNA obtained from NPAF or NF, exons 1 through 6 of the SMPD1 gene were PCR-amplified using exon-specific primers. Sequence analysis was performed using routine procedures for molecular diagnostics of Niemann-Pick-type A disease, i.e. dye terminator based sequencing reactions with purified amplification products from the above mentioned PCR-reactions, separation of products by capillary electrophoresis on a Beckmann CEQ8000 instrument and sequence analysis with the aid of *Codon Code Aligner* software.

### ASM-assay

The activity of ASM in cell lysates was measured using an ELISA-kit from Echelon (K3200) following the protocol provided by the supplier. In brief, cells cultured in T75 flasks, were harvested, washed, re-suspended in 0.5ml of 0.9% NaCl with 1mM PMSF, and subjected to four freeze/thaw cycles in liquid nitrogen. Following determination of protein concentration using a NanoDrop instrument (Thermo Fisher Scientific), cell lysates and standards were processed according to the protocols supplied, and incubated for 3h at 37°C. The reaction was stopped and fluorescence (excitation 360nm, emission 460nm) was read in a SpectraMax iD3 instrument (Molecular Devices).

### ATP-assay

Cellular ATP levels were measured using a luminometric luciferase assay of as described elsewhere (Haugwitz et al., 2006). In brief, cells were seeded at a density of 2×10^4^ per well of a 96-well tissue culture plate. The next day, samples were treated as detailed in figure legends and text, and cellular ATP content was measured using the ATP Bioluminescence Assay Kit CLSII from Roche and a Lumat 9705 instrument from Berthold.

### β-hexosaminidase assay

The β-hexosaminidase release assay using 4-methylumbelliferyl N-acetyl-ß-D-glucosaminide as a substrate was as described previously (Rodriguez et al., 1997). In brief, cells were grown in 12-well tissue culture plates. After removal of media, cells were washed with in HBSS with HEPES and Ca^2+^, and α-toxin or SLO in HBSS with HEPES and Ca^2+^ was added. Controls received buffer only. Fluorescence was measured in a TriStar LB 941 instrument from Berthold Technologies equipped with appropriate filters.

### Flow cytometry

For analysis of annexin V-binding and propidium iodine (PI) influx, cells were treated or not with α-toxin as indicated in the figures, washed twice with PBS, detached with trypsin, re-suspended in complete culture medium, and washed twice with ice cold PBS. The pellet was re-suspended in 100 μl annexin-binding-buffer (10mM HEPES, 140mM NaCl, 2.5mM CaCl_2_ pH 7.4), with 1μl PI stock solution (100μg/ml) plus 3.5μl annexin V Alexa Fluor^TM^ 647. After incubation for 15 min at RT 400μl of annexin-binding buffer was added and samples were kept on ice. Samples were analyzed using a BD FACS CANTO II and FACS Diva software.

### Monitoring changes in [Ca^2+^]_i_

To detect potential changes in [Ca^2+^]_i_ in response to α-toxin we used an AM-ester of the calcium-sensitive fluorescent dye Fluo-8, which is cleaved by cellular esterases. We employed two assay formats: a bulk-assay in micro-titer plates and video-microscopy for evaluation at the single cell level. PFT-induced changes of [Ca^2+^]_i_ in cells were monitored by using Fluo-8AM. Cells (2-3 x 10^4^/well in 200μl complete medium) were seeded in a 96-well format using black flat-bottom microplates (Greiner) and incubated at 37°C o/n. Cells were loaded for 30min with a mixture of Fluo-8AM (100μM), Cremophor, 0.1% (w/v) and probenecid, 2mM. Loaded cells were washed, toxin was added and fluorescence intensity recorded (60s intervals, up to 20min) in a SpectraMax iD3 instrument (excitation 485nm, emission 535nm).

For analysis by video-microscopy, cells were seeded into channels of IBIDI µ-slides VI ^0.4treat^ at a density of 5×10^4^ cells per well. The next day, cells were incubated with a freshly prepared solution containing 5μM Fluo-8AM, 0.1% cremophor and 2mM probenecid in cell culture medium for 30min at 37°C. Subsequently, cells were washed with HBSS and live cell imaging was performed in both fluorescence and phase contrast mode using an inverted widefield microscope (Leica Thunder Imager) equipped with an environmental control box (37°C and 5% CO_2_) and a 40x/0.95 air objective. Cells were imaged directly in the channels of the IBIDI µ-slides under a constant flow of fresh HBSS or toxin dilution ensured by a pump. Cells were recorded with HBSS flow, typically for 10 - 50 frames, before HBSS was replaced by pore forming toxins diluted in HBSS and recorded for at least another 110 frames at 5 s intervals.

### Monitoring changes of plasma membrane potential

Changes of PM potential were measured by using DiSBAC2(3), Thermo Fisher Scientific (B418), a slow response membrane potential sensitive dye. The day before the experiment cells were seeded at a density 3×10^4^ cells per well in black 96-well plates (Greiner). After 2 times washing with HBSS/Hepes/Ca^2+^, DiSBAC2(3) dissolved in ultrapure DMSO was added to give a final concentration of 100nM of the dye. After an initial 7min equilibration-period, toxins were added to cultures and fluorescence recordings (excitation 525nm, emission 565nm) continued for up to 40min at RT.

### Fluorescence microscopy

For indirect IF experiments, cells were grown on glass coverslips O/N and incubated or not with toxins and/or inhibitors as detailed in the figure legends. After incubation with primary antibodies, cells were washed in PBS, fixed in 2% paraformaldehyde in PBS for 10min at RT and incubated with AlexaFluor®-488-conjugated secondary antibodies. To investigate whether cells internalize α-toxin we treated them with Alexa546-labelled α-toxin, fixed, and either treated them with 0.1% Triton-X100 for 10min at RT, to permeabilize the PM, or left them un-permeabilized. Subsequently, samples were blocked with 3% BSA in PBS, and Alexa546-labelled α-toxin was immuno-stained with rabbit-anti-α-toxin F(ab)_2_ and Alexa 488-conjugated secondary antibody. For the simultaneous detection of internalized Alexa546-labelled α-toxin cells and caveolin-1, Alexa546-α-toxin-treated cells were fixed and permeabilized as described above, and caveolin was stained by indirect IF using anti-caveolin-1 antibody N20 and Alexa 488-conjugated secondary antibody. For visualization of macro-pinosomes and α-toxin in context, cells were serum-starved for 24h before adding FITC-dextran 70kDa (1mg/ml) with or w/o Alexa546-labelled α-toxin (250nM) and incubated at 37°C for 15min. Subsequently, coverslips were gently washed and fixed (3% paraformaldehyde, 15min at RT). Coverslips with stained cells were mounted on slides with ProLong^TM^ Diamond Antifade Mountant. Samples were examined with a Zeiss Axiovert 200M epifluorescence microscope equipped with a Plan Apochromat 100x/1.4 aperture lens. Digital images were acquired with a Zeiss Axiocam using Zeiss AxioVision software (4.1).

### Statistics

Numerical data displayed in graphs represent mean values ± SEM from n ≥ 3 independent experiments, unless stated otherwise. Statistical significance of differences between mean values was generally assessed using one-way ANOVA and Tukeýs post hoc test, if not stated otherwise. To reveal trends of some kinetic data we used simple linear regression. For selection and application of tests for significance we employed GraphPadPrism8; for details see figure legends. Significance was generally assumed at *P* <0.05.

## Acknowledgements

We like to thank PD Dr. med. H. Rossmann, Department of Laboratory Medicine University Medical Center Mainz, for help with sequence analysis. We gratefully acknowledge technical assistance by J. Altmeier from FACSlab, Institute for Toxicology, University Medical Center Mainz.

## Author contributions

GvH and MH conceived of the study. Claudia N and Carolin N, MM, GvH and SR performed experiments, acquired and analyzed data. GvH and MH designed experiments, interpreted data and wrote the manuscript. All authors contributed to manuscript revision, read and approved the submitted version.

## Abbreviations

AMPK: AMP activated protein kinase
ASM: acid sphingomyelinase
CDC: cholesterol dependent cytolysins
CIDRE: Ca^2+^ influx-dependent repair
CYD: cytochalasin D
DISBAC2(3): Bis-(1,3-Diethylthiobarbituric Acid)
EGFR: epidermal growth factor receptor
ESCRT: endosomal sorting complex required for transport
FACS: fluorescence activated cell sorting
GCN2: general control non-derepressable 2
HBSS: Hank’s balanced salt solution
ILV: intraluminal vesicles
NF: normal fibroblasts
NPAF: Niemann-Pick type A fibroblasts
O/N: overnight
PFP: pore forming proteins
PFT: pore forming toxins
PhlyP: phobalysin P
PI: propidium iodide
PM: plasma membrane
PPP1r15B: protein phosphatase 1 regulatory subunit 15B
PS: phosphatidylserine
SLO: streptolysin O
SLO-MBP: streptolysin O-maltose binding protein
SMPD1: sphingomyelin phosphodiesterase 1 gene
SREBP: sterol regulatory element binding protein
TRP: triparanol
VCC: *Vibrio cholerae* cytolysin

