## supplementary Figures for "Recovery of fibroblasts from membrane attack by *S. aureus* α-toxin does not depend on acid sphingomyelinase but involves macropinocytosis": suppl Figures von Hoven et al bioRxiv 12082023.pdf

**Supplementary Figure 1**

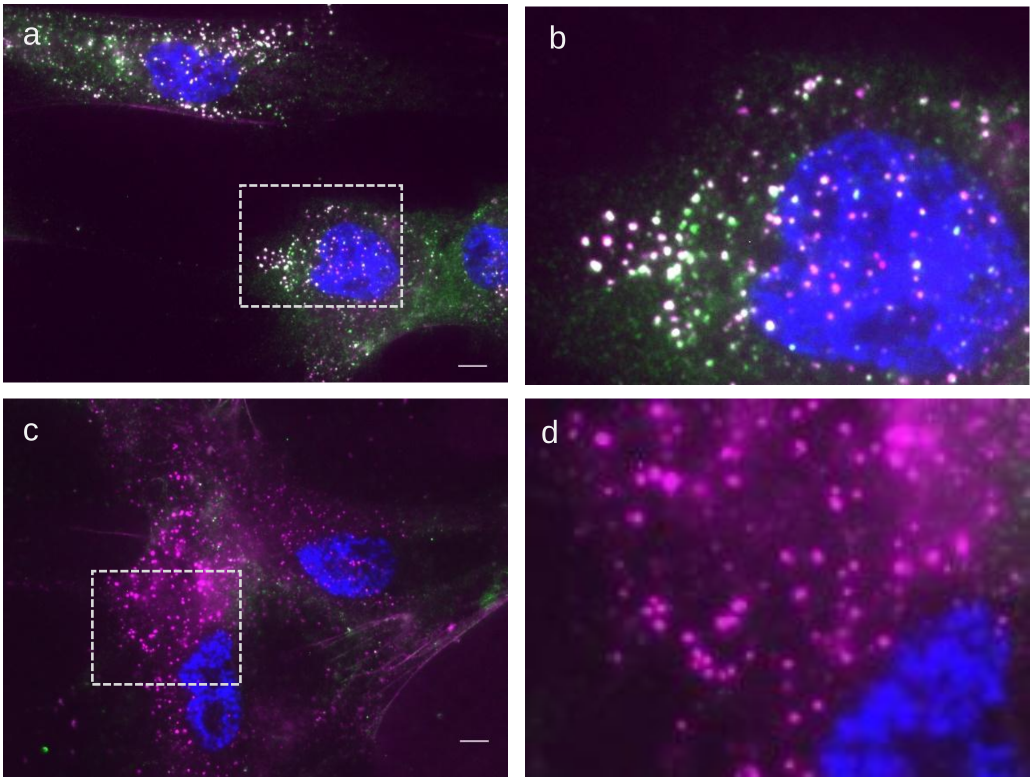


***S. aureus α*-toxin enters fibroblasts lacking functional ASM**. NPAF grown on coverslips were incubated with Alexa546-α-toxin, 250nM (~ 8 µg/ml), for 15min, Samples were fixed and either permeabilized with 0.1% Triton (a,b), or not (c, d), before immunostaining with rabbit anti-α-toxin F(ab)2 and Alexa 488-conjugated secondary antibody. Samples were analysed by wide-field IF-microscopy. Representative images. Size bar, 5µm.

**Supplementary Figure 2**

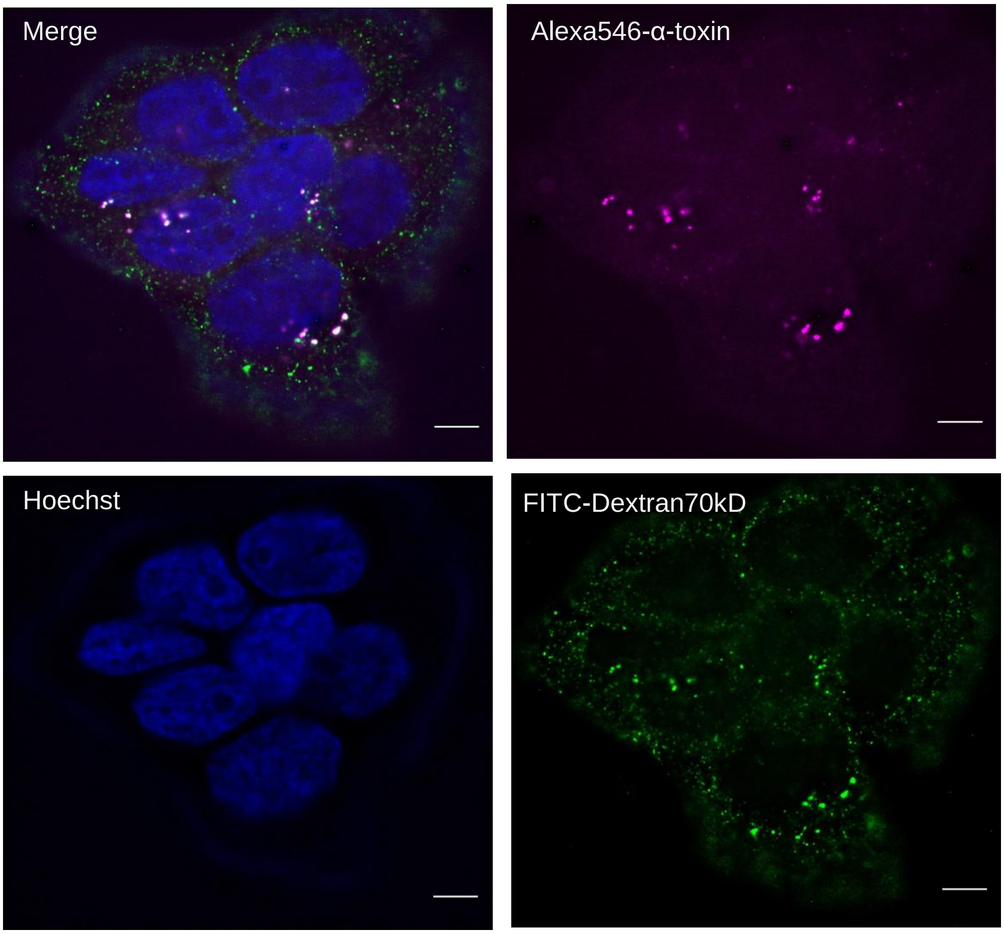


***S. aureus* α-toxin co-localizes with FITC-Dextran70kD in HaCaT cells.** HaCaT cells were incubated for 15min with Alexa546-α-toxin (2μg/ml) and FITC-Dextran70kDa (1mg/ml). Samples were analysed by wide-field IF-microscopy. Representative image. Size bar, 5 μm.
